# Exercise exacerbates decline in the musculature of an animal model of Duchenne muscular dystrophy

**DOI:** 10.1101/360388

**Authors:** KJ Hughes, A Rodriguez, A Schuler, B Rodemoyer, L Barickman, K Cuciarone, A Kullman, C Lim, N Gutta, S Vemuri, V Andriulis, D Niswonger, AG Vidal-Gadea

## Abstract

Duchenne muscular dystrophy (DMD) is a genetic disorder caused by loss of the protein dystrophin. In humans, DMD has early onset, causes developmental delays, muscle necrosis, loss of ambulation, and early death. Current animal models have been challenged by their inability to model the early onset and severity of the disease. Thus it remains unresolved if increased sarcoplasmic calcium observed in dystrophic muscles follows or leads the mechanical insults caused by the muscle’s disrupted contractile machinery. This knowledge has important applications for patients, as potential physiotherapeutic treatments may either help or exacerbate symptoms, depending on how dystrophic muscles differ from healthy ones. Recently we showed how burrowing dystrophic (*dys-1*) *C. elegans* recapitulate many salient phenotypes of DMD, including loss of mobility and muscle necrosis. Here we report *dys-1* worms display early pathogenesis, including dysregulated sarcoplasmic calcium, and increased lethality. Sarcoplasmic calcium dysregulation in *dys-1* worms precedes overt structural phenotypes (e.g. mitochondrial, and contractile machinery damage) and can be mitigated by silencing calmodulin expression. To learn how dystrophic musculature responds to altered physical activity, we cultivated *dys-1* animals in environments requiring high amplitude, or high frequency of muscle exertion during locomotion. We find that several muscular parameters (such as size) improve with increased activity. However, longevity in dystrophic animals was negatively associated with muscular exertion, regardless of the duration of the effort. The high degree of phenotypic conservation between dystrophic worms and humans provides a unique opportunity to gain insights into the etiology of the disease, as well as the initial assessment of potential treatment strategies.

**SIGNIFICANCE:** Duchenne muscular dystrophy is a degenerative disease affecting tens of thousands of people in the US alone. Much remains unknown about the disease, including the chain of events that links the loss of dystrophin to muscle death, or the extent to which exercise might be able to protect degenerating muscles. We used the nematode *C. elegans* to show that sarcoplasmic calcium dysregulation takes place in dystrophic muscles long before other overt signs of damage manifest. When placed in assays that altered muscular activity by increasing either contraction frequency or amplitude, we observed several metrics associated with muscular repair increase. However, no treatment positively affected the life expectancy of dystrophic animals.

## INTRODUCTION

Duchenne muscular dystrophy (DMD) is a lethal muscle-wasting disease affecting 1 in 3500 males. It is characterized by progressive muscle weakness, loss of ambulation, and premature death^1, 2^. To date there is no cure for Duchenne muscular dystrophy. DMD is caused by loss-of-function mutations in the gene encoding the dystrophin protein, resulting in its absence from muscles^3-5^. At the sarcolemma, dystrophin links cytoskeleton to the extracellular matrix through a complex of membrane proteins, where it helps protect myocytes from the powerful forces generated by their contractile machinery. Additionally, dystrophin participates in nitric oxide (NO) signaling cascade^6-8^. Absence of dystrophin may cause muscle shearing and increased sarcoplasmic calcium levels, potentially leading to muscle decline directly through cytotoxicity^9^, mitochondrial damage^10^, or additional mechanisms^11^.

DMD patients display dramatic levels of muscle degeneration, and lose mobility early on in life^1^. Progress in DMD is hindered by lack of genetic model systems that accurately recapitulate the severe loss of mobility and muscle degeneration seen in humans. Similar deficits are only observed in canine models^12, 13^ where many factors prevent interrogation of the molecular basis of the disease^14^. More (genetically) amenable organisms such as flies, worms, and mice have not matched the phenotypic severity observed in humans or dogs^15-17^. However, progress resulted from matching dystrophin mutations with sensitizing mutations in other related proteins (or through other insults), as these animals more closely recapitulate the acute motor and muscular decline seen in humans^18^. These approaches are useful, but also cloud interpretation of their results as it becomes challenging to unequivocally assign a phenotype to loss of dystrophin. Lack of phenotypic severity in these systems may result from compensatory mechanisms (e.g. utrophin upregulation^14, 19^), and/or the shorter lifespan of these animal models. Alternatively, their attenuated phenotypes may result from insufficient muscular challenge to animals that are (mostly) kept under low exertion regimens not matching those they would experience in their natural environment. Consistent with this idea, muscle degeneration was directly correlated with strength of muscular contraction in (mdx) mice modeling DMD^20^. This suggests that modeling the disease accurately requires appropriate behavioral paradigms, capable of modulating muscle exertion in behaving animals.

One aspect of DMD that remains unanswered is the sequence of molecular steps that lead from loss of dystrophin to muscle cell death. Increases in sarcoplasmic calcium have been reported in dystrophic animals^21^, as have the differential expression of genes involved in calcium homeostasis^22^. However, what causes calcium levels to increase in the first place, or which altered gene expression profiles are the result of compensatory (vs dysregulated) mechanisms remains unknown. At a more practical level, it is not known the extent to which muscular exertion may be beneficial or detrimental in the treatment of DMD^23, 24^. Specifically, there is no consensus regarding the potential benefits of physical therapy to treat DMD. Therefore, it remains unclear if there is a type and level of exercise that might prove prophylactic for dystrophic muscles, or if muscle activity and decline are irrevocably linked in this disease^25, 26^. Answering these questions has direct and immediate implications for the tens of thousands patients currently enduring this disease^27^. Faced with these limitations, a recent roundtable session convened at the New Directions in Biology and Disease of Skeletal Muscle Conference. Several recommendations were agreed upon to improve understanding of the role of muscular exertion in the etiology of DMD^25^. Chief among these recommendations was the development of new assays and animal models better reflecting the disease in humans. This would permit the use of genetic techniques to understand the etiology of DMD, as well as to identify and evaluate therapeutic approaches.

We developed a burrowing assay that permits the modulation of substrate density and the resulting muscular exertion produced by burrowing animals^28^. Under this paradigm, animals modeling DMD genetically recapitulated the acute motor and muscular decline seen in DMD patients^29^. Here we show that burrowing in dystrophic *C. elegans* the early onset and sarcoplasmic calcium dysregulation characteristic of DMD. To evaluate the potential impact of different physical activities on dystrophic musculature we harnessed distinct natural behaviors performed by worms (crawling, burrowing, and swimming). This allowed us to modulate the force, amplitude, or frequency of muscular exertion by dystrophic animals. We report that sarcoplasmic calcium increase occurs before onset of other overt muscular phenotypes and that, consistent with in vitro work in mice, calcium release appears spared while calcium reuptake is challenged in the muscles of freely behaving dystrophic animals. Furthermore, calcium reuptake is restored by silencing calmodulin expression. Modulating the duration and intensity of exercise treatments we find that the longevity in dystrophic animals is inversely related to the muscular exertion produced by these animals, but not to the frequency at which the animals performed these activities.

## METHODS

### Strains

Animals were cultured on nematode growth media (NGM) agar plates and fed *Escherichia coli* OP50. Plates were held at 20°C as previously described^30^. *Caenorhabditis elegans* strains wild-type N2, BZ33 *(dys-1(eg33))*, LS292 *(dys-1(cx18))*, and ZW495 (zwIs132 [*myo-3p::GCamp2 + lin-15(+)*]) were obtained from the Caenorhabditis Genetic Center. The Caenorhabditis Genetics Center (CGC) is supported by the National Institutes of Health Office of Research Infrastructure Programs (P40 OD010440). HKK5 and HKK22 strains (kind gift from Hongkyun Kim) are BZ33 and N2 strains (respectively) expressing nuclear and mitochondrial GFP in body wall muscles, *ccIs4251[Pmyo-3GFP-NLS, Pmyo-3 GFP-mit]* (Oh & Kim 2013). The AVG6 strain was obtained by crossing the ZW495 and BZ33 strains.

### RNA interference

To silence expression of dystrophin *(dys-1)*, calmodulin *(cmd-1)*, sarcoplasmic reticulum (SR) calcium pump (SERCA, encoded by *sca-1)*, calsequestrin *(csq-1)*, in WT or dystrophic animals, we cultivated worms (N2 or BZ33) in bacteria containing RNAi control vector (L4440), or a vector targeting the gene of interest as previously described^31^.

### RNA extraction and quantitative PCR

Brightness differences between dystrophic and wild-type worms was assessed by expressing the calcium reporter GCaMP2 using the promoter for myosin-3 *(Pmyo-3::GCaMP2)*. To assess if differences in brightness between wild-type and dystrophic animals reflected differences in *myo-3* expression between these strains we measured myosin levels using qPCR following the protocol described by Laranjeiro et al. (2017)^32^. Because the greatest difference between the strains was observed in the L1 stage, we collected about 200 wild-type and dystrophic L1 larvae (ZW495 and AVG6) into TRIzol reagent (Thermo Fisher Scientific, Waltham, MA, USA) and froze animals in liquid nitrogen. Total RNA was then extracted following directions from manufacturer. We used Quantitect reverse transcription kit (Qiagen, MD, USA) to create a total cDNA from RNA and remove genomic DNA. Removal of genomic DNA was confirmed by PCR amplification (using cDNA) of genes spanning intronic regions.

We conducted quantitative PCR using diluted cDNA, PowerUp SYBR green Master Mix (Thermo Fisher), and 0.5μΜ gene specific primers in a QuantStudio^tm^ 5 Real Time PCR System (Thermo Fisher). We calculated relative myosin expression in ZW495 and AVG6 L1 worms using the ΔΔCt method^33^ using the cdc-42 as a reference gene^34^.

### Imaging

#### Confocal Imaging

We used a Zeiss LSM 510 Confocal laser scanning microscope to acquire 10x or 100x images. Muscle mitochondria and nuclei were imaged directly by placing animal in mounting media containing 10% 100mM sodium azide and immediately acquiring the images.

#### Filming

We used a Flea2 camera (Point Grey, Vancouver) mounted on an Olympus SZX12 stereomicroscope to acquire sequences of (1032×776 pixels) TIFF images. Swimming and crawling animals were filmed at 15 and 7.5Hz (respectively) for 1 min, but burrowing animals were filmed for 5 minutes at 1.8Hz (owing to the slow velocity of this behavior).

#### Image analysis

We used Image-Pro-7 (Media Cybernetics, USA) to detect and track worm centroids as previously described^35^. IgorPro (Wave Metrics) was used to analyze neck bending amplitudes.

#### Calcium transients

All calcium comparisons were conducted on animals filmed under identical illumination and magnification conditions (alternating strains during the same filming session). Calcium signals were recorded with the aid of an X-cite 120 PC Q light source. We used ImageJ to measure GCaMP2 signals. We measured the brightness of the head (seventh and eighth muscles in each quadrant), midbody (opposite the vulva: sixteenth muscles), and the tail of each worm (twenty third and twenty fourth muscles). Figure Supp 2A shows a schematic of how images were analyzed. We measured differences in maximum brightness between antagonist contracting and relaxing muscles, as a segment reached the apex of contraction. Each calcium measurement was accompanied with a body curvature measurement corresponding to the bending of the segment under consideration. The strength of muscle contraction is reported as the ratio of maximum brightness of the contracting to the relaxing muscles.

***Calcium dynamic changes*** over time were obtained by measuring the maximal brightness of the dorsal and ventral muscles posterior to the terminal pharyngeal bulb (as described above) for three to five consecutive crawling cycles. Calcium changes were plotted as maximum brightness of the eighth body wall muscles over time. To compare the kinetics of calcium movement in the cytosol we normalized calcium signals to their range and cycle duration (Figure 3 and Figure Supp 2F).

#### Calcium imaging vs animal velocity

To compare the velocity of freely moving animals with the brightness of their muscles we filmed crawling worms as described above, using only blue fluorescent light to illuminate animals and stimulate GCAMP2 expression. Magnification was decreased in order to produce a filming arena of about 1cm^2^. Animal velocities were then obtained using ImagePro as described above. Importantly, we report maximal (whole) animal brightness for this set of experiments as the reduced magnification made single muscle measurements unfeasible.

### Behavioral Assays

#### Continuous burrowing

To test the effect of muscle exertion on worms we placed animals in agar volumes (200μL) of different densities (1%, *3%*, or 6%) containing dissolved OP50 *E. coli* for nourishment. Animals were prevented from exiting the burrowing volume by surrounding them it in every direction with 8% chemotaxis agar, which worms are normally unable to penetrate. The shallowness of the test agar (~1mm) allowed us to constrain the animals within a narrow vertical range, and facilitated filming (Figure Supp 1).

#### Intermittent burrowing

The effects of intermittent burrowing were assessed by allowing the animals to burrow for 90 minutes every day for five days. After a session, animals were allowed to crawl onto a petri plate seeded with OP50 where they crawled between burrowing sessions.

***Swimming velocity*** was assessed as previously described^35^. Briefly five to ten (never-starved) day-one adult animals were placed in a 1 cm diameter copper ring on an agar plate. The ring was flooded with 250 μL liquid NGM, which closely mimics the chemical makeup of their culture plates and the internal osmolarity of worms. Animals were filmed after acclimating for two minutes.

#### Continuous swimming

15 adult worms were placed in 10 cm NGM agar plates and flooded with 4 ml of liquid NGM containing OP50 bacteria in suspension as food source. After three days, animals were transferred to new plates to prevent inclusion of F1 adults in our measurements.

#### Intermittent swimming

This was conducted as described above. After 90 minutes swimming, animals were transferred to an OP50 bacterial lawn to crawl until the following day. Treatments continued daily for five days.

***Crawling*** was filmed and assessed on day one adult animals (or after 1, 3 and 5 days for the exercise experiments). Care was taken that animals were clean of OP50 bacteria by rinsing them in liquid nematode growth media (NGM). To control for animal handling, crawling animals that were compared to intermittent swimming or burrowing animals were transferred the same number of times.

***Larval behavior treatments*** were similar to those described above for adults. Recently (~1hr) hatched L1 larvae were filmed to obtain crawling velocity, size, and brightness. Alternatively, they were transferred into a 1μl drop of liquid NGM, and injected into burrowing pipettes, or placed in either swimming or crawling plates. After three days incubation at 20°C (standard time to adulthood for crawling WT worms), animals were allowed to crawl out of the agar and assessed.

#### Longevity experiments

The longevity experiments involved subjecting animals to continuous or intermittent swimming, burrowing, and crawling until all animals had died or could no longer be accounted for. We performed at least five replicates of each behavioral assay with fifteen day one adults. During the experiments, offspring were separated from adult worms to ensure they were not being included in longevity analysis. This was accomplished by transferring adults to new assay environments every three days until they ceased laying eggs.

#### Immunohistochemistry

To image muscular f-actin we used a protocol adapted from Ono (2001)^36^. Worms were transferred to a fixative solution containing PBS and 16% formaldehyde, which was then replaced with acetone, and placed in -20°C for 5 minutes to permeabilize the cuticles. Animals were then rinsed with PBS-TG, which we replaced with 1:500μL of dried down Phalloidin and rotated overnight. The worms were then rinsed with PBS-TG and stored at 4°C until mounted in mounting media.

#### Statistics

We used SigmaPlot (Systat Software, inc.) to assess differences in velocity between conditions, including Kruskal-Wallis one-way ANOVA on ranks with Dunn’s Method for multiple comparison as a follow up. Calcium imaging was analyzed using standard one-way ANOVA’s and the Holm-Sidak multiple comparisons follow up test. Regression lines for WT and dystrophic calcium release and reuptake kinetics were compared using ANCOVA. Longevity was evaluated using Kaplan-Meier estimate, allowing us to include censored data (animals that were unable to be accounted for; cannot assume dead or alive, Kaplan and Meier (1958)^37^. Cox proportional hazard was used to assess survival when comparing the effect of both type of exercise *and* duration of exercise^38^. To see summary and comparative statistics ran in this study please refer to Supplementary Table 1.

## RESULTS

### Burrowing dystrophic worms recapitulate many of the features of Duchenne muscular dystrophy

One of the earliest diagnostic features of DMD is impaired, and eventual loss of, ambulation^39, 40^. Day-1 adult worms of two independently derived dystrophic alleles, *dys-1(eg33)* and *dys-1(cx18)*, show severe burrowing impairments when burrowing in 3% agar (Fig. 1A). To determine if the motor impairments observed were related to muscle health, we obtained WT and dystrophic animals with GFP localized to the body wall musculature and mitochondria (kind gift by Dr. Hongkyun Kim). Muscle degeneration and necrosis are characteristics of DMD that we similarly saw recapitulated in dystrophic worms. After burrowing in 3% agar for five days, adult dystrophic worms develop swollen, bursting, or absent mitochondria and cell nuclei in their musculature (Fig. 1B). Furthermore, burrowing dystrophic worms *dys-1(eg33)* had a reduced lifespan compared to healthy WT animals (Fig. 1C). Burrowing dystrophic *C. elegans* worms recapitulate some of the most salient phenotypes of the disease in humans. They model Duchenne muscular dystrophy genetically, in motor and muscular degeneration, and in reduced longevity, without the need for sensitizing mutations. In humans, loss of function mutation in dystrophin manifest early-on during childhood^41^. To determine if burrowing larval *C. elegans* recapitulated the progression of the disease in humans we turned to test developing dystrophic worms.

**Figure 1.**
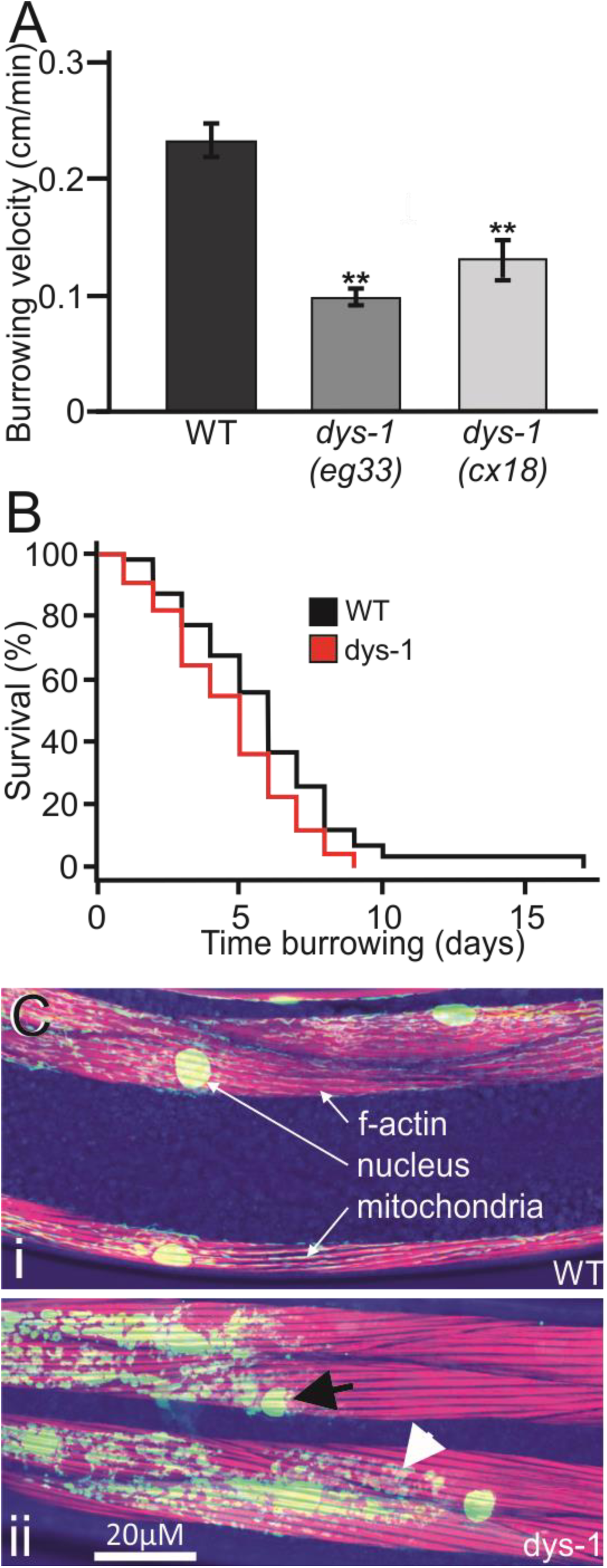
Burrowing *dys-1* mutants recapitulate cellular, behavioral, and lifespan phenotypes associated with Duchenne muscular dystrophy. A. *dys-1* mutants carrying independent loss-of-function alleles *eg33* and *cx18*, show decreased velocity when burrowing through 3% agar. > 5 assays/ condition; > 50 animals/ condition; Holm-Sidak comparisons. **B.** In comparison with wild-type animals that burrowed for five days (i), dystrophic worms (ii) showed fewer mitochondria, and nuclei. Red is phalloidin staining of f-actin, green is GFP localized to nuclei and mitochondria. **C.** Compared to wild-type animals, adult dystrophic worms cultivated in 3% agar showed decreased longevity. > 8 assays/ condition; > 160 animals/ condition; Cox proportional hazard. **p<0.01.

### Developing dystrophic animals show enhanced deficits that parallel the disorder in humans

Cytosolic calcium dysregulation has been proposed as one possible mechanism responsible for the decline of dystrophic musculature. To measure and compare calcium levels in the musculature of dystrophic and healthy animals we crossed the dystrophic BZ33 strain (*dys-1(eg33)*) with the ZW495 strain [*myo-3p::GCamp2 + lin-15(+)*], which expresses the calcium reporter GCaMP2 in the body wall musculature of wild-type animals under the control of the myosin-3 promoter. The result was the generation of the AVG6 strain which expresses GCaMP2 in the body wall muscles of dystrophic *(dys-1(eg33))* animals. Recently hatched dystrophic L1 larvae had significantly lower crawling velocities than wild-type L1 larvae (*p=0.018, Figure 2A). Additionally, compared to wild-type larvae, dystrophic animals were significantly brighter (Fig. 2B, p<0.001). We were surprised by the observation of increased sarcoplasmic calcium levels in recently hatched animals, as these had little time to damage their musculature by mechanical means. Nevertheless, these animals showed the head hyper curvature characteristic of dystrophic adults^42^. Because the strain we used employed myosin to drive GCaMP2 expression, it was possible that dystrophic worms might just express greater levels of myosin-3 leading to more GCaMP2 and brighter signals. To assess this possibility we extracted RNA from wild-type and dystrophic L1 larvae (as previously described^32^) and measured relative myosin-3 levels against a housekeeping gene previously reported to show stable expression during development *(cdc-42)34*. We found that the ΔΔCt for myosin-3 was not significantly different in the two strains (0.07 fold change). We therefore concluded from these experiments that differences in muscle brightness reflect differences in calcium levels rather than differences in GCaMP2 reporter levels.

**Figure 2.**
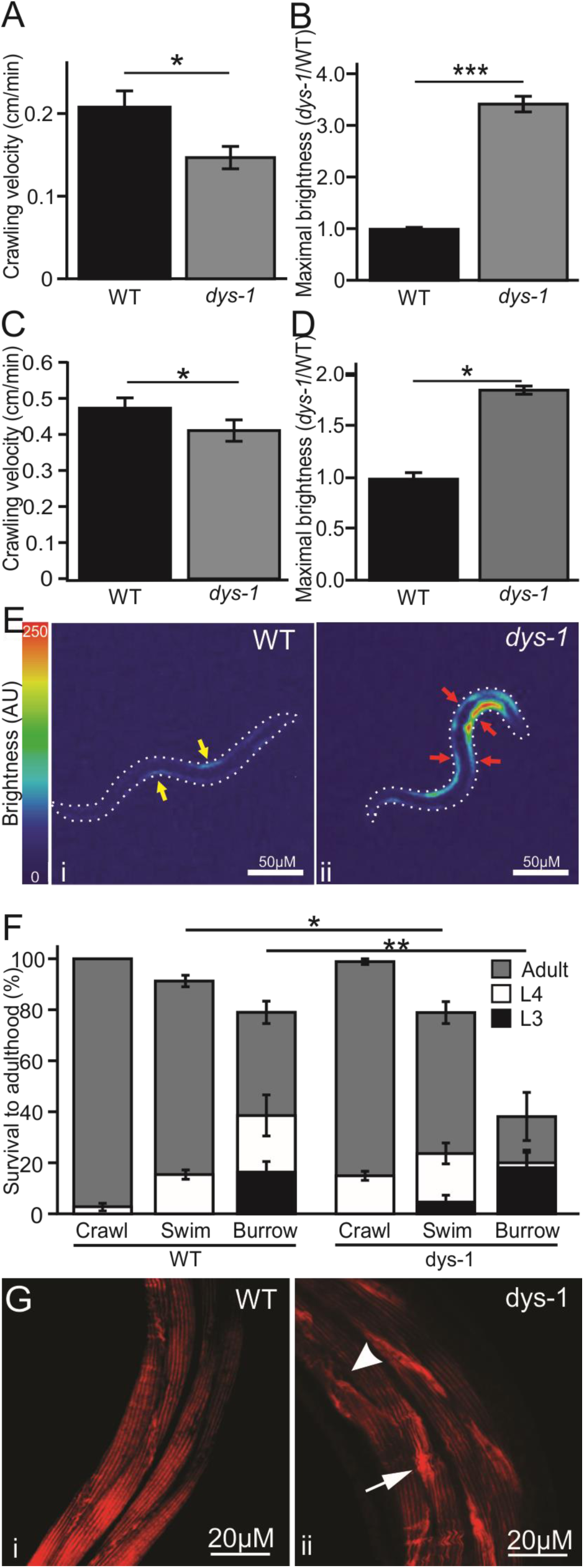
Dystrophic *C. elegans* display motor and physiological deficits early in development. **A.** Recently hatched dystrophic L1 larvae have reduced crawling velocity compared to wild-types. > 10 animals/ condition; t-test. **B.** Muscular cytosolic calcium concentration was measured indirectly through GCaMP2 expression in body wall musculature (driven by the myosin promoter, *Pmyo-3::GCaMP2*). Young dystrophic larvae (L1) are more than three times brighter than wild-types. Maximal (whole animal) brightness is normalized to wild-type. > 20 animals/ condition; Mann-Whitney Rank Sum Test. **C.** Heat map showing the relative brightness of wild-type L1 larvae (i), and dystrophic L1 larvae (ii, AU = arbitrary units). Wild-type animals show low amplitude signals in the contracting musculature and no signal in the opposite (relaxed) muscles (yellow arrows). Dystrophic larvae display the characteristic hypercontracted shape, brighter calcium signals, and show evidence of concomitant activation of antagonistic muscles (red arrows). D. Compared to wild-type, or to other locomotion treatments, burrowing *dys-1* larvae showed increased mortality and developmental delays with roughly half of surviving animals reaching adulthood after three days. **E.** After burrowing in 3% agar until adulthood for three days, now adults, dystrophic animals continued to have locomotion impairments and increased cytosolic calcium levels **(F). G.** Dystrophic animals that did survive burrowing to adulthood had signs of muscular degeneration. ***p<0.001, **p<0.01, *p<0.05.

In healthy worms, antagonistic muscle groups alternate their contraction cycles to generate rhythmic locomotion. Accordingly, in wild-type larvae only contracting muscles showed GCaMP2 activation (yellow arrows in F2Ci). However, in dystrophic larvae elevated calcium was detected through GCaMP2 in both sets of antagonistic muscles (Fig 2.Cii). Inability to timely, and fully, clear sarcoplasmic calcium during the relaxation part of the cycle might necessitate ever greater calcium transients from contracting muscles to achieve locomotion.

To evaluate the effect that differential muscular exertion had on dystrophic larvae, we cultivated L1 worms to adulthood (t=3 days) in 3% agar, liquid growth media (NGML), or crawling on petri plates. After three days incubation at 20°C, we found that surviving dystrophic worms had greater developmental delays than wild-type regardless of treatment (Fig. 2D). However, compared to wild-types and other treatments, burrowing produced the greatest developmental delays, with only half of surviving dystrophic worms reaching adulthood by this time point. In addition to developmental delays, burrowing dystrophic larvae also suffered increased mortality. Only 40% of dystrophic animals were alive after three days incubation in 3% agar. Similar results were obtained when the agar density was increased to 6% (Fig. Supp 3C). In contrast, wild-type larvae had twice (80%) that survival rate. High mortality and developmental delays have both been associated with Duchenne muscular dystrophy^43, 44^.

After burrowing in 3% agar to adulthood, we evaluated dystrophic (now) adult worms for locomotion, sarcoplasmic calcium transients, and cellular integrity. Dystrophic adults maintained their (pre-burrowing) larval trend, with significantly lower than wild-type velocity (Fig. 2E), and higher sarcoplasmic calcium levels (Fig. 2F). We noted that both of these parameters were now closer to wild-type levels. However, the musculature of dystrophic worms that burrowed from larvae to adulthood displayed signs of degeneration. Dystrophic muscles had greater accumulation of f-actin, and signs of fibre tearing rarely found in their wild-type counterparts (Figure 2G). Dystrophic *C. elegans* thus appear to match the human disease not only in loss of motility and muscular degeneration, but also in terms of its early onset, developmental delays, and high mortality.

While informative, their small size (~250μm) and transient nature make larvae less suited to dissect out the interaction between different parameters affecting the progression of the disease. To investigate the interaction between contractile machinery integrity, sarcoplasmic calcium, and motor function, we turned once again to adult animals. The maximal brightness of the calcium signal recorded from moving adult animals correlated with their moving velocity, and with their body curvature (Fig. Supp. 2B, C). Similar to L1 larvae, dystrophic worms displayed a similar (but larger) range of curvatures, but significantly larger maximal brightness in their neck muscles (Fig. Supp. 2D,F). We next turned to evaluate how muscle activity and output changes over time in healthy and dystrophic animals.

### Dystrophic worms display increased levels of sarcoplasmic calcium

To assess the progression of calcium dysregulation, we evaluated the brightness of adult animals as they burrowed in 3% agar over time. Day-1 adults were placed in burrowing wells and evaluated after one, three, and five days. We found that both wild-type and dystrophic animals had a similar contracted to relaxed brightness ratios, indicating similar fold changes in sarcoplasmic calcium during contractions. This ratio decreased over time, but remained similar between the two strains (Fig. 3A). Unlike wild-type animals, dystrophic worms experienced a drop in burrowing velocity after the third day (Fig. 3B). To try to understand how motor output (measured as velocity) might become uncoupled from calcium transients (i.e. brightness) we looked closer at the minimal and maximal levels of sarcoplasmic calcium during contraction and relaxation in freely burrowing wild-type (Fig. 3C) and dystrophic (Fig. 3D) animals. We found that in wild-type animals, maximal sarcoplasmic levels of calcium remained constant over time for actively contracting muscles. However, as the animal ages, the calcium levels in the antagonistic (relaxed) muscles increase. This change might explain the drop in the ratio of contracted-to-relaxed brightness measured in Fig. 3A. Turning to look at the dystrophic animals, we saw a different picture. While the ratio of contracted-to-relaxed muscle tracked that of wild-types over time (Fig. 3A), the cytosolic concentration of calcium in these animals was different. We found that the antagonistic (relaxed) muscles in dystrophic animals had elevated levels of sarcoplasmic calcium that remained constant over time. Actively-contracting muscles also showed elevated (above wild-type) levels of sarcoplasmic calcium, which decreased steadily over time. By day five, this drop caused an overlap in the levels of sarcoplasmic calcium in both contracting and relaxing muscles, helping to explain the loss of propulsion observed at this time point. The relative stable burrowing velocity, and contracted to relaxed calcium levels suggests that animals endeavor to meet some external demands necessary to achieve propulsion.

**Figure 3.**
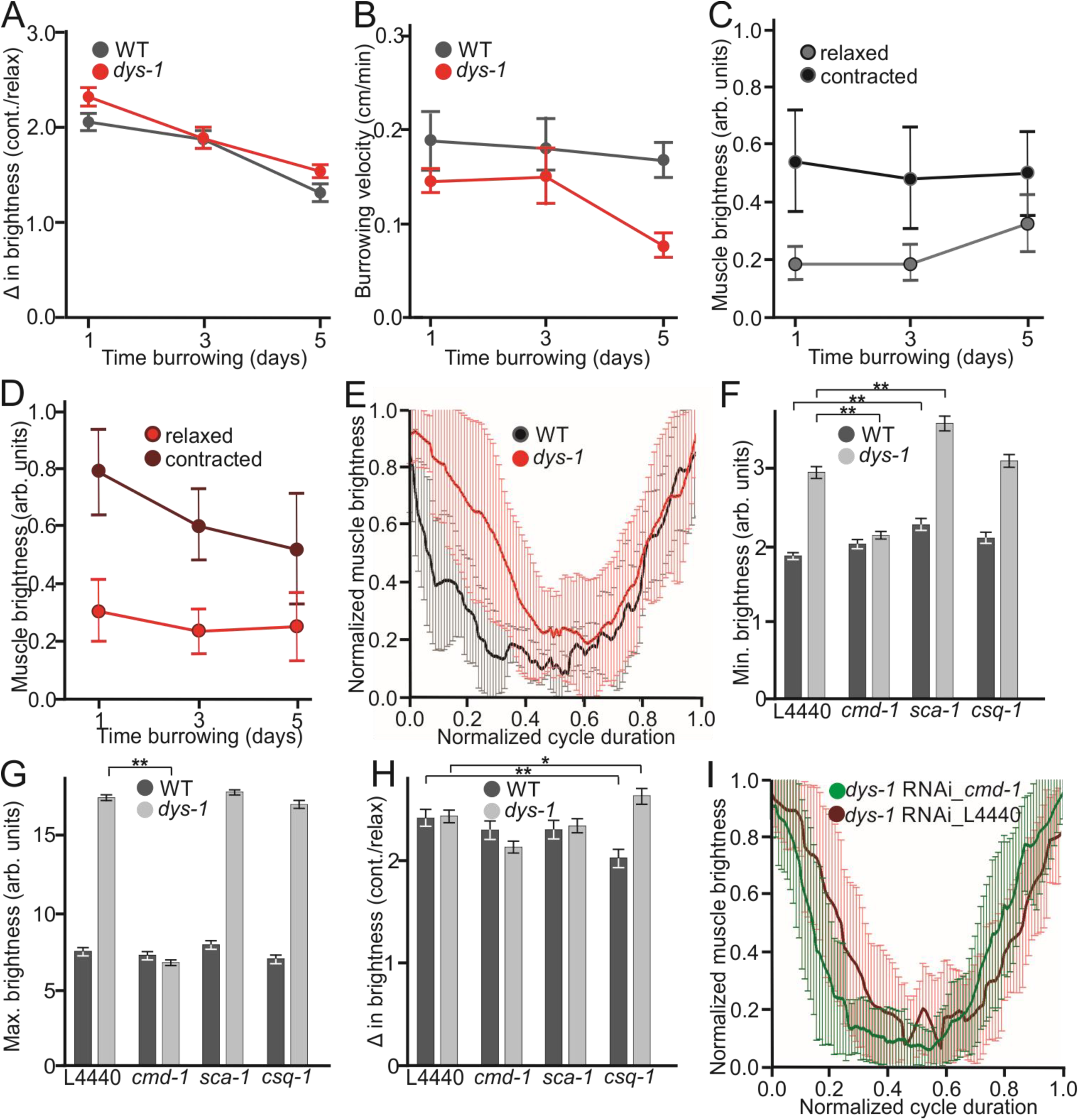
Calcium release becomes uncoupled from muscle contraction in dystrophic musculature. **A.** Wild-type and dystrophic worms burrowing continuously in 3% agar over their first five days of adulthood. Calcium transients associated with muscular contractions are displayed as the ratio of maximal GCaMP2 brightness of contracting over relaxing (contralateral) muscles at the peak of contraction. The initial calcium transient ratio (contracted/relaxed), and their subsequent decline with age, is similar for wild-type and dystrophic animals. **B.** Velocities of wild-type and dystrophic animals burrowing in 3% agar overtime. By day 5 of burrowing, dystrophic animals show a decline in burrowing velocity compared to wild-types. **C.** Ranges of maximal cytosolic calcium for dorsal/ventral (contracting/ relaxing) muscles over time. Wild-type animals show low contracted, and basal calcium levels throughout most of the experiment. **D.** Dystrophic animals maintained elevated basal levels of cytosolic calcium in relaxed musculature, as well as greater levels in contracting muscles which decline over time. **E.** Kinetics of calcium transients during muscle contraction evaluated indirectly through GCaMP2 brightness measurements. Muscle brightness was measured during several (>5) crawling cycles in (>5) wild-type, and dystrophic animals. Maximal brightness is shown normalized to cycle duration, and muscle brightness is normalized to entire brightness range. Dystrophic worms show delayed and incomplete calcium reuptake during the relaxation phase of the cycle (0.0-0.5), but show no difference from wild-types during the contraction phase of the cycle (0.5-1.0). We used RNAi to silence calmodulin *(cmd-1)*, SERCA *(sca-1)*, and calsequestrin *(csq-1)* in wild-type and dystrophic worms. **F.** Silencing SERCA expression increased basal calcium levels in both dystrophic and wild-type animals. Silencing calmodulin expression retuned dystrophic animals to wild-type calcium levels for their basal brightness and maximal brightness during contraction **(G). H.** Silencing calsequestrin significantly affected amplitude of the calcium transient during contractions in distinct manners, increasing the ratio for dystrophic, and decreasing the ratio for wild-type worms. **I.** Kinetics of calcium transients for dystrophic worms fed empty RNAi vectors (L4440), or vectors targeting calmodulin expression *(cmd-1)*. *p<0.05, **p<0.01.

### Higher basal levels of sarcoplasmic calcium are associated with delays in reuptake, leading to incomplete relaxation

Dysregulation in cytosolic calcium levels has been reported in dystrophic musculature^45^. Does this dysregulation affect every calcium process during muscle contraction, or are there events that are particularly susceptible to this insult? To investigate this, we quantified the dynamics of calcium cycling during muscular contractions. We filmed and analyzed GCaMP2 signals from the dorsal and ventral muscles behind the pharynx of wild-type and dystrophic worms (muscles 7 and 8) as they freely crawled on agar plates (Fig. 3E). To compare calcium kinetics across strains and animals, we normalized the timing to one movement cycle (starting at the angular peak for the body segment under consideration (cycle time = 0), progressing through the peak of relaxation (cycle time =0.5), and returning back to the peak of the next body wave (cycle time = 1). We next normalized the data by the range of muscle brightness. These transformations allowed us to compare the timing of calcium changes, minimizing concern for movement velocity and signal magnitude. Comparisons of the cycling of sarcoplasmic calcium between wild-type and dystrophic animals showed that both strains had similar calcium release kinetics during the contraction phase (Fig. 3E, second half of the plot). However, dystrophic worms had significantly faster, but incomplete, calcium reuptake during the relaxation phase (Fig. 3E, first half of the plot, and quantified in Fig. Supp 2F). The incomplete clearance of sarcoplasmic calcium during the relaxation phase of muscle contractile cycle is consistent with the increased levels of calcium measured for burrowing dystrophic worms (Fig. 3D), and the increased brightness of both larval and adult dystrophic animals (Figs. 2B,C,F, and 3C). These findings also are in agreement with in vitro calcium recordings from dystrophic mice fibers, where dystrophic muscle fibers were shown to display altered calcium reuptake but normal calcium release^46^.

### Silencing calmodulin in dystrophic muscles restores calcium levels and kinetics

To further investigate the nature of intracellular calcium movement in wild-type and dystrophic animals we used RNA interference (RNAi) to silence the cytosolic calcium binding protein calmodulin *(cmd-* 1), SERCA, which is responsible for moving calcium into the SR *(sca-1)*, and calsequestrin *(csq-1)* which is responsible for calcium buffering inside the SR. We found that silencing the worm homolog of SERCA (SCA-1) significantly increased the basal levels of calcium in both dystrophic and wild-type animals (Fig. 3F). Surprisingly, only silencing calmodulin was able to reduce the minimal and maximal brightness of dystrophic muscles down to wild-type levels (while not affecting wild-type animals, Fig. 3F,G). Consistent with its proposed role helping regulate calcium release during contraction, silencing calsequestrin only significantly affected the change in brightness during contraction (Fig. 3H). Previous work suggested that increases in cytosolic calmodulin associated with dystrophic muscles might inactivate SERCA^47^. This could explain how silencing cytoplasmic calmodulin helps restore calcium levels by SERCA inactibation. To investigate this possibility, we compared the kinetics of calcium movement in dystrophic animals where calmodulin had been silenced, versus animals fed the empty RNAi vector (L4440). We found that silencing calmodulin in dystrophic animals restored their rate of calcium reuptake back to wild-type levels (Fig. 3I). Our results suggest that previously reported increases in cytosolic calcium levels in dystrophic muscles^11, 48^ might lead to increased cytosolic calmodulin^49^, and inactivation of SERCA.

Whether muscle decline is ushered by impaired calcium trafficking, or by mechanical damage from its contractile machinery has important implications for the type of muscle, and activities, that might be most susceptible in dystrophic patients. If the contractile machinery is largely responsible for muscle decline, it might be possible to recruit the muscular repair response by means of low amplitude/high frequency exercise (e.g. swimming) without compromising integrity in the process. However if calcium dysregulation plays an important role in the etiology of DMD, high frequency exercise might lead to increased rates of calcium release and reuptake and overwhelm dystrophic muscles even with decreased contraction amplitude. To evaluate the potential benefit of differential muscle activation in dystrophic muscles we turned to the natural behavioral repertoire of worms.

### Locomotion through different physical environments places distinct demands of the musculature of *C. elegans*

One important unanswered question Duchenne muscular dystrophy patients face is what is the likelihood and extent to which treatments based on physical activity might be beneficial to dystrophic musculature. Evidence from different studies has been inconclusive^50^. We therefore set out to test if *C. elegans* might be useful to determine the potential impact of different activity regimens on dystrophic musculature.

***C. elegans*** crawl on solid surfaces, swim in liquids, and burrow through dense media^35, 29, 51^. During these types of locomotions, worms adopt distinct S, C, or W body shapes (Fig. 4A). Each of these forms of locomotion has distinct kinematics. Swimming is characterized by the fastest body bend frequencies, and the lowest bending amplitudes. Crawling has larger body bend amplitudes, and intermediate frequencies. Burrowing worms produce the greatest bending amplitudes, at the lowest frequencies. In addition, the force output of burrowing worms can be modulated by varying the density of the media through which they move (Fig. 4B). Swimming is the fastest of the three behaviors, and burrowing the slowest (Fig. 4C). The anterior end of worms houses a greater number of muscles than medial or posterior regions of similar length. Not surprisingly, the head of *C. elegans* appears to contribute to propulsion disproportionately compared to the rest of its body (Fig. 4D). This asymmetry is reduced for burrowing animals, where the posterior musculature appears to be recruited more equitably in order to achieve locomotion (Fig. 4E).

**Figure 4.**
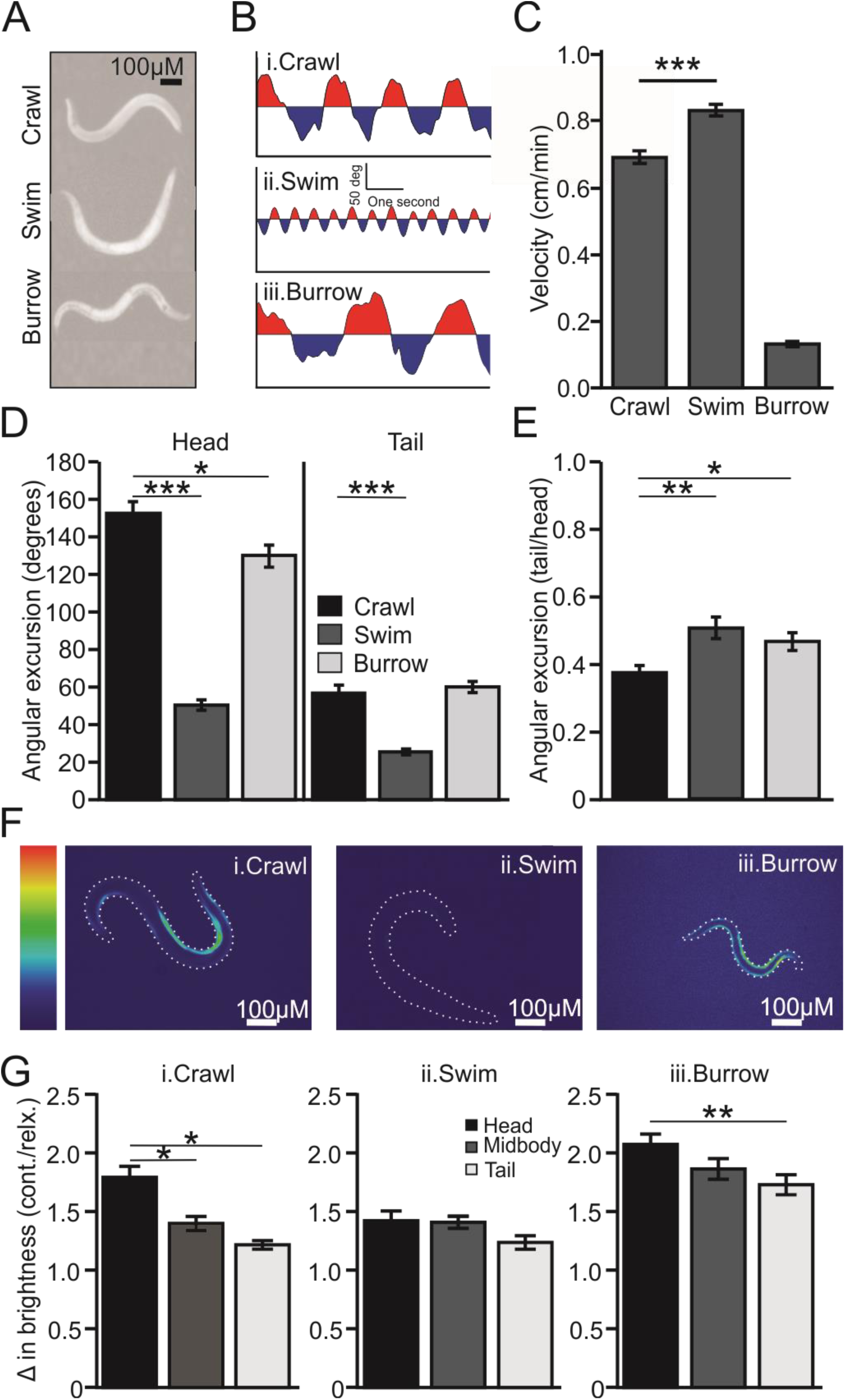
Locomotion through different environments is associated with distinct outputs from the musculature of *C. elegans* **A.** *C. elegans* crawls on solid surfaces, swims in liquid, and burrows through media adopting characteristic “S”, “C”, and “W” shapes respectively. **B.** Neck curvature plots for crawling, swimming, and burrowing worms illustrating that burrowing consists of low frequency and high amplitude movements, while swimming relies on high frequency and low amplitude movements. **C.** Each of these behaviors has characteristic velocity ranges. Swimming worms move the fastest, and burrowing worms the slowest. **D.** Movement amplitudes differs significantly between behaviors, and across the length of the animals. Animals crawl using large head flexions that do not propagate fully to their tail. **E.** During swimming and burrowing, the contribution of the tail of the animals to locomotion showed a significant increase compared to crawling. **F.** The anterior bias in propulsion for all behaviors, as well as the increasing contribution of posterior musculature during swimming and burrowing are reflected by muscular calcium transients. Heat map showing variations in cytosolic calcium in body wall musculature inferred from GCaMP2 fluorescence (driven by the myosin-3 promoter). Anterior is right for all animals. **G.** Quantification of calcium transients for different areas of the body during crawling, swimming, and burrowing. While slower, burrowing produced the greatest calcium transients across the entire body of the worm. ***p<0.001, **p<0.01, *p<0.05.

To determine if the different kinematics observed within body regions and between behaviors was correlated with muscular exertion, we filmed animals expressing GCaMP2 in their musculature (driven by the *myo-3* promoter, Fig. 4F). Paralleling our kinematic measurements, burrowing animals displayed the strongest change in brightness. While still displaying posterior dampening, burrowing animals produced the greatest calcium transients across their entire bodies. In contrast, the musculature of swimming worms produced calcium transients that were lower in magnitude to crawling or burrowing, but similar in amplitude across the length of their bodies. Therefore swimming in *C. elegans* requires fast and shallow movements across the whole body that don’t require great muscular output. In contrast, burrowing consists of slow, deep body contractions that require powerful output from the animal’s musculature. We conclude that these distinct locomotor modalities can be used to mimic regimens involving high frequency and low amplitude (swimming), or high amplitude and low frequency (burrowing) activity.

### Dystrophic muscles do not benefit from increased frequency or amplitude of contraction

Next we set out to determine if changes in the motor demands placed on dystrophic musculature had the potential to delay or prevent some of the phenotypes associated with muscular dystrophy. We placed dystrophic day-1 adult worms to either crawl in agar plates, to burrow in 3% agar wells, or to swim in liquid NGM. We measured their velocity and calcium transients after one, three, and five days of treatment. As shown in Fig. 5A and B, burrowing worms displayed a continuous decline in velocity mirrored by a decrease in the brightness ratio of contracted-to-relaxed musculature (Fig. 5B). In contrast, swimming and crawling animals maintained their velocity until day three, with swimming worms showing the steepest loss in mobility by day 5. These observations were matched by the calcium measurements in these treatments. Swimming worms continuously increased the magnitude of their calcium transients between day one and five, while crawling worms increased this metric only after the third day of treatment (Fig. 5B).

**Figure 5.**
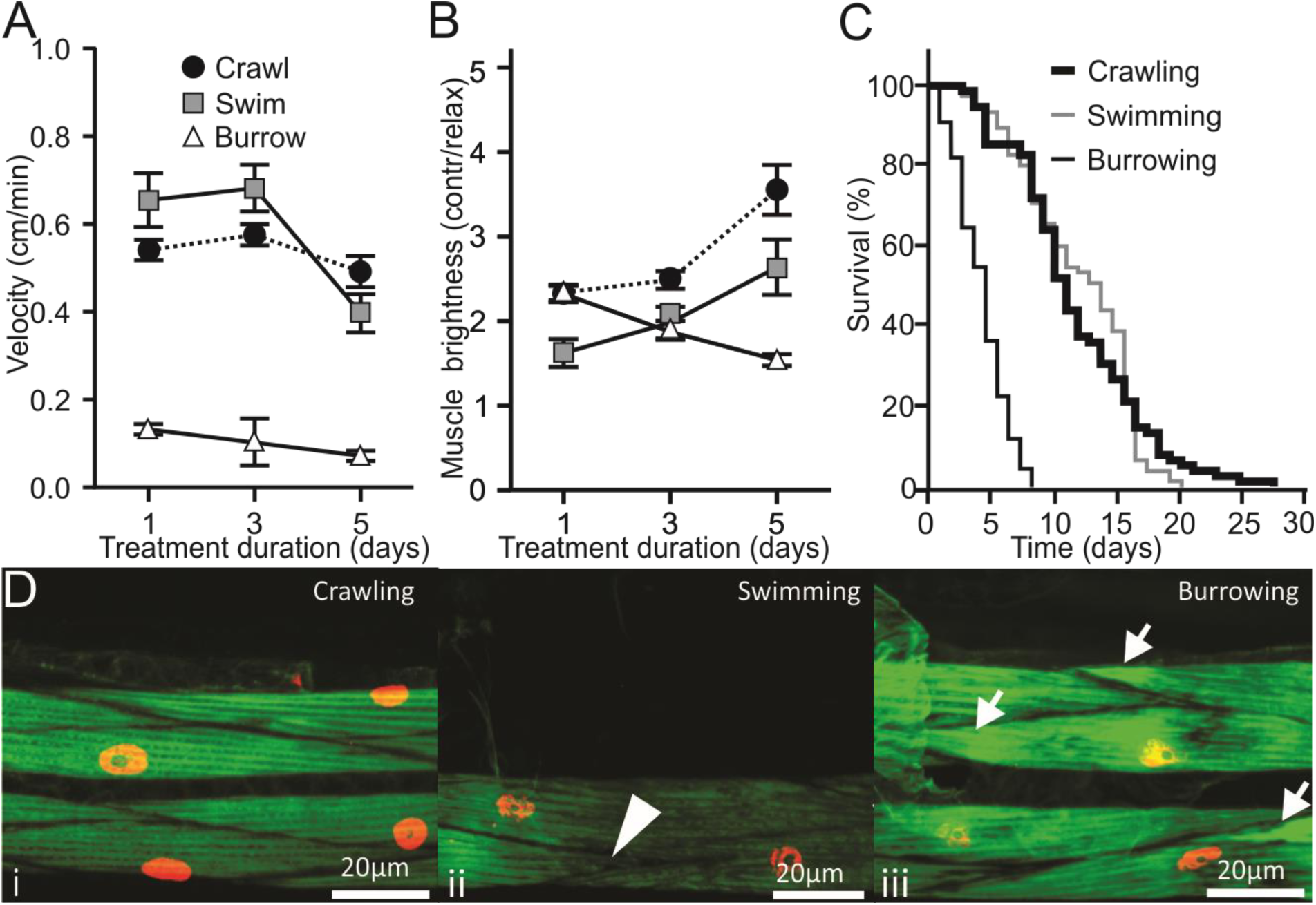
Physical exertion negatively impacts health and longevity in dystrophic *C. elegans* worms. **A.** The velocity of burrowing animals declined continuously throughout the experiment, while animals maintained under swimming or crawling treatments maintained their velocities until day 3. **B.** The decline seen by day 5 crawling and swimming animals was accompanied by a similar increase in the magnitude of calcium transients. **C.** While swimming and crawling differ in velocity and in calcium transient amplitudes, both activities had similar effects on dystrophic worm longevity. However, burrowing decreased animal longevity significantly compared to other treatments (Cox proportional hazard, vs. crawling: p<0.001; vs. swimming: p<0.001) **D.** Representative confocal images of dystrophic muscles following crawling, swimming, or burrowing for five days. Swimming and burrowing animals had signs of degeneration, including torn or fewer actin fibers in swimming animals (arrows), and f-actin accumulation at the attachment plaques of burrowing animals (arrowheads).

These data suggest that dystrophic worms faced with a burrowing challenge are able to increase the release of sarcoplasmic calcium during contraction in order to achieve propulsion through solid media, even when faced with the additional challenge of overcoming the increased basal activation of antagonistic musculature. However, this ability declines continuously, as they experienced a drop in calcium transients and burrowing velocity throughout the duration of the experiment (Fig. 5A,B). In contrast to burrowing, the demands placed on dystrophic musculature by swimming seem milder. Animals were able to continuously increase the amplitude of their calcium transients and maintain constant output until the third day of treatment. After this point, calcium transients became uncoupled from motor output (Fig. 5A, B). As expected, crawling appeared to be the least challenging of the three treatments. Crawling animals maintained motor output throughout the duration of the experiment, only needing to increase the amplitude of calcium transients after the third day. For every treatment, maintaining motor output became more challenging over time. Animals were either able to continuously increase effort and maintain output (swimming), or lost their ability to generate propulsion (burrowing). By the end of the burrowing treatment (day 5), only about 50% of the animals were still alive (Fig. 5C). Burrowing animals had significantly decreased longevity compared to crawling (p<0.001) and swimming (p<0.001). However we observed no significant difference in longevity between crawling and swimming (p = 0.354). Confocal images of the body wall musculature following treatment showed that the f-actin fibers of crawling worms were mostly intact (Fig. 5Di). Swimming worms had wider muscle cells with lower density of actin fibers, that showed signs of damage (Fig. 5Dii). Burrowing worms showed the greatest signs of muscle damage, with evident tears in actin filaments, and areas of increased actin depositions (Fig. 5Diii).

Swimming and burrowing are activities that differ in their muscular demands. Burrowing requires high power generation, while swimming requires high contraction frequency. While neither of these activities increased longevity or muscle health (as measured by ultrastructure, output, and calcium dynamics), burrowing had a clear negative effect on every metric considered. The experiments performed above consisted of chronic treatments where animals were continuously required (in order to achieve locomotion) to propel themselves through the prescribed media. It is possible however that such treatment might overwhelm the coping ability of dystrophic muscles, and that a more limited treatment might produce instead beneficial effects when chronic treatment would do the opposite. To evaluate this possibility we repeated the experiments above but reduced their duration to ninety minutes of daily activity. This duration was selected based on a recent study by the Driscoll lab showing that 90 minutes of swimming by *C. elegans* were sufficient to trigger gene expression associated with exercise in humans^32^.

### Altering exercise duration affected several muscular parameters but not animal longevity

From our previous experiment we found that continuous burrowing had profoundly negative effects on dystrophic musculature, and that continuous swimming did not improve the animal’s outcomes above crawling levels. To determine if reducing the duration of these activities might produce a beneficial effect we reduced treatments to 90 daily minutes. Animals were placed in crawling plates between treatments where they were free to crawl.

Dystrophic animals swimming 90 minutes per day had a reduced drop in swimming velocity compared to worms continuously swimming (Fig. 6A). However, like continuously swimming worms, these animals continued to increase calcium transients over the five days of the experiments. Similar to swimming, worms that burrowed 90 minutes per day showed increased velocity compared to those that burrowed continuously (Fig. 6C). However, the greatest impact on velocity was restricted to the first day of burrowing. By the third day, these burrowing worms displayed the same rate of decline in velocity and calcium transient amplitude as those burrowing continuously (Fig. 6C, D).

We were surprised to see that burrowing for 7.5 hrs over five days (intermittent burrowing treatment) had similar effects on dystrophic animals than burrowing 120 hrs over the same period of time (continuous burrowing). To evaluate the effect of burrowing duration on dystrophic muscles we used phalloidin to image f-actin fibers in the musculature of these animals. We found that by the fifth day of treatment, dystrophic worms that burrowed continuously showed clear signs of actin fiber damage, including tears in the filaments and areas of increased accumulation (Fig. 6 E). In contrast, worms that burrowed 90 min per day did not show actin accumulation. They did however have areas where actin fibers were stressed. When comparing these treatments, we noticed that the size of muscle cells differed between treatments. Worms that burrowed once per day had significantly larger muscle cells and were themselves significantly larger than worms that burrowed continuously (Fig. 6 F). This effect was even greater for healthy wild-type worms, which were about twice as large as wild-type worms burrowing continuously. These improved muscle metrics led us to predict that worms burrowing 90 minutes daily might show increased longevity compared to those burrowing continuously. This was however not the case (Fig. 6G). Despite improvements in velocity, and muscle cell growth, longevity was not improved for animals that burrowed once daily. Similar effects were measured for swimming animals (Fig. Supp. 3).

## DISCUSSION

### Modeling Duchenne muscular dystrophy with nematodes

The understanding of the molecular mechanisms responsible for muscle degeneration and death in Duchenne muscular dystrophy has benefited from the many animal models ranging from mice, to dogs, zebrafish, flies, and worms^14^. Each system has intrinsic advantages and limitations, which explains their continued ability to contribute to the understanding of this disorder.

One limitation plaguing all animal models of DMD (except dogs) is the lack of severity of the disorder in these organisms. Therefore, most studies have resorted to a combinatorial approach combining dystrophin loss with loss in an associated protein, or with other type of sensitization^15,52,53^. The nature of these approaches complicates interpretation of their findings, and draws attention to the obvious question of why humans show such an acute phenotype compared to other animals? The limitations of current animal models created gaps in understanding that coalesced during the roundtable session convened at the New Directions in Biology and Disease of Skeletal Muscle Conference^25^. Several recommendations were agreed upon to improve our understanding of the role of muscular exertion in the etiology of DMD. Chief among these was the development of new assays in animal models to reflect the physical limitations observed in patients, as this would permit the use of genetic techniques to identify new therapeutic targets. These and other needs were recently formalized in the 2015’s Action Plan for the Muscular Dystrophies developed by the NIH and the Muscular Dystrophy Coordinating Committee (MDCC)^54^.

To answer these stated needs, we developed a new assay that allows us to use the genetically tractable nematode *C. elegans* to investigate the etiology of DMD and identify promising strategies for its treatment. Our burrowing assay produces phenotypes that approach those seen in DMD^29^. We found that burrowing dystrophic worms track many of the phenotypes associated with DMD in humans. These include increased cytosolic calcium, early onset, loss of mobility, mitochondrial damage, contractile machinery failure, muscle death, and shortened life span. To our knowledge, this is one of the most complete replications of the human prognosis documented in an animal model of DMD.

### Impaired reuptake of cytosolic calcium associated with increase in basal calcium levels, and increased calcium release during muscle contractions

We found that recently hatched L1 dystrophic larvae increased levels of calcium compared to wild-type animals (Fig. 2B, C). These animals had limited opportunity to move, and showed no overt signs of muscle degeneration. The data suggest an initial uncoupling of calcium dysregulation from physical exertion and supports the idea that increased cytosolic calcium is (at least initially) not necessarily the result of sarcolemmal damage caused by the absence of dystrophin^9, 55^. However increased basal levels of cytosolic calcium arise, they seem to remain high throughout the life of the dystrophic animal (Fig. 3D). In contrast, wild-type animals only see their basal calcium increase to dystrophic levels by the fifth day of burrowing.

Increased basal levels of calcium could result in muscles being in a continuous state of contraction, as calcium would remain in the cytosol to activate the contractile machinery. This inappropriate tension would in turn increase the work required from the appropriately contracting (antagonistic) muscles during their normal activation. We found that burrowing dystrophic worms increase their calcium transient enough to produce the same ratio of contracted-to-relaxed muscle measured in wild-type animals (roughly a twofold increase in brightness for contracted over relaxed muscles). Maintaining this ratio seems important to maintain velocity during locomotion. Thus animals lose mobility when this ratio drops, due either to decreases in calcium release during contraction (e.g. dystrophic animals), or to increases in basal levels (e.g. aging wild-type animals).

Because dystrophic animals seem to be capable of (at least initially) appropriately upregulate calcium release and maintain muscular output, calcium dysregulation does not seem to be the consequence of a global collapse in calcium cycling. Supporting this idea, we found that the kinetics of calcium release during contraction remain mostly unaffected in dystrophic animals. However, this was not the case for calcium reuptake during the relaxation phase of contraction. This was significantly accelerated in dystrophic animals, however the large amount of calcium present in the cytosol prevented this modulation from accomplishing the complete clearance of calcium in time for the beginning of the next contraction. Work on mice reported similar increases in calcium reuptake (but not in release) in mice modeling muscular dystrophy^46^. In this study, Rittler reported an upregulation in the expression of genes involved in calcium reuptake into the sarcoplasmic reticulum of dystrophic animals (such as the SERCA1 and 2). Previous work also pointed to increased levels of calmodulin as responsible for the inhibition of SERCA 1^56^. Our findings point to a conserved mechanism for early calcium dysregulation during the onset of the disease (prior to overt dystrophic phenotypes in dystrophic worms or mice). Supporting this idea we found that silencing calmodulin restored levels and kinetics of cytosolic calcium (Fig. 3F-I). Together with previous work, our data suggest a mechanism for the maintenance of elevated calcium levels in dystrophic muscles whereby an initial increase in cytosolic calcium is countered by increases in calmodulin and in SERCA expression. However elevated levels of calmodulin inactivate the SERCA pumps and prevent calcium from being reuptaken into the sarcoplasmic reticulum. Understanding the mechanism by which basal cytosolic calcium increases in dystrophic musculature remains of paramount importance, as restoring normal calcium levels without compromising the integrity of cellular organelles could remove one of the factors contributing to the accelerated decline of dystrophic musculature.

### Harnessing natural behaviors to study the role of exertion in dystrophic muscle health

One of the important aspects of Duchenne muscular dystrophy that remain unanswered is the extent to which muscular exertion might be beneficial or detrimental in the treatment of DMD^23^. To address this need we used the natural behaviors performed by worms and evaluated their differential loads on the muscular system. Worms crawl on surfaces, swim through liquids, and burrow through solid media^19, 35^. Most *C. elegans* research consists of studying animals crawling on an agar plate. Crawling consists of high amplitude body movements of intermediate frequency (~0.5Hz). *C. elegans* crawl using mostly their (more numerous) anterior musculature, and appear to find this less challenging than other types of locomotion. In contrast with crawling, swimming is a fast behavior (~2HZ) and consists of high frequency body bends of low amplitude. Recent work from the Driscoll lab^32^ demonstrated that a single swimming bout in *C. elegans* could induce key features reminiscing of exercise in mammals. In their natural habitat, *C. elegans* likely spend most of their time burrowing through solid decaying plant matter or soil. Burrowing consists of slow, low frequency movements of high amplitude. Therefore, comparing these three locomotor strategies in worms it is possible to test the effect of activities of high frequency and low amplitude (i.e. swimming), and high amplitude and low frequency (i.e. burrowing). Furthermore, by varying the density of the burrowing media, it is possible to modulate the force required to achieve locomotion^19^. Our calcium measurements from animals moving through these media support the idea that burrowing requires the greatest muscular exertion as contracting muscles were twice as bright as antagonistic (relaxing) muscles. In contrast, swimming and crawling required contractions only about 50% brighter than the relaxing baseline (Fig. 4F, G).

### Dystrophic animals required progressively greater calcium transients to maintain motor output

We found that over time, even swimming and crawling dystrophic worms had to increase the amplitude of calcium transients in their musculature to maintain their velocity. This seems to support the idea that as dystrophic damage takes its toll in the musculature, greater effort must be made to achieve the same propulsion. Paradoxically, this greater effort would result in further damage to the muscles, creating a runaway effect and eventually uncoupling calcium transients from motor output once damage has progressed sufficiently. Indeed, we found that by day five both crawling and swimming worms increased their calcium transients dramatically while also experiencing their greatest loss in propulsion (Fig. 5A, B). The ability of worms to modulate calcium release to offset ongoing damage was limited and depended on the magnitude of the task. For example, burrowing worms started with large calcium transients which quickly decreased in amplitude as the musculature became compromised. These animals saw a continuous decrease in velocity throughout the experiment. While swimming had no effect on dystrophic worm longevity, it did lead to tearing of acting fibers in the musculature of dystrophic animals (Fig. 5C, D). Burrowing on the other hand had profound negative effects on longevity and muscle actin integrity. We conclude that neither chronic swimming, nor chronic burrowing are able to improve the health of dystrophic muscles or the longevity of animals modeling the disease.

### Longevity of dystrophic animals is determined by the intensity of the exercise performed, and not by its duration

Since Driscoll’s group detected exercise-like responses in *C. elegans* muscles after only 90 minutes swimming^32^, we reasoned that reducing the time animals spent in each exercise treatment might improve their prognosis. We therefore repeated all our experiments but this time animals in the swimming or burrowing treatments were allowed to perform these activities for ninety minutes per day. The remainder of the time animals were placed back in crawling assay plates (intermittent treatment). We found that dystrophic animals that swam 90 minutes daily showed improvements over those that swam continuously in terms of their velocity and calcium transient amplitudes (Fig. 6A, B). Worms that burrowed 90 minutes per day showed improvement over those burrowing continuously only during the first day of treatment. By the third day of treatment both velocity and calcium transients had experienced significant drops (Fig. 6C, D). Examining the structure of the muscles we found that actin fibers showed signs of degeneration that increased the longer the animals were allowed to burrow (Fig. 6E). Surprisingly, the muscle cells of animals that burrowed intermittently appeared larger than those of worms burrowing continuously (Fig. 6Eii vs iv). We found that not only were continuously burrowing dystrophic worms larger than their wild-type counterparts, worms that burrowed intermittently were significantly larger than worms that burrowed continuously (Fig. 6F). This finding suggests that intermittent burrowing in *C. elegans* triggers muscle hypertrophy characteristic of resistive training in humans. The greatest hypertrophic change was recorded for wild-type animals which had twice the area of worms that burrowed continuously. Combining ours, with the results on swimming by Laranjeiro et al^32^, we suggest that swimming and burrowing in *C. elegans* might be investigated for their ability to mimic endurance and resistance exercise (respectively). Unfortunately, unlike wild-type animals where longevity correlated with the time spent performing physical exertion, the longevity of dystrophic worms was determined by the maximal exertion performed, rather than by its duration Fig. Supp. 3).

If conserved, these results suggest that endurance and strength exercise treatments have the potential to improve some muscle metrics such as muscle size, or output, while also hastening muscle fiber degeneration, and significantly reducing longevity. The contractile apparatus of dystrophic animals (a primary target of exercise treatments) is not necessarily tied to the fate of the cell, or the animal. Of the potential intracellular targets for mitigating degeneration in dystrophic muscles, releasing the inhibition of the SERCA pump by high levels of calmodulin appears to be the most promising approach for reducing overall cytosolic calcium levels and slow down the mechanical damage caused by the contractile machinery in the absence of dystrophin.

Much work remains to be done to understand how loss of dystrophin leads to loss of muscle and death. Great efforts are currently underway to harness gene or transcription editing with the goal of restoring dystrophin levels in muscles. While important and promising, these treatments will have limited applicability to the tens of thousands of patients suffering with this disease today. Therefore, complementary to those efforts, it is important to devise ways to improve the health of those currently suffering with DMD. This will require understanding the etiology of the disorder, and devising treatments that improve quality of life without compromising life expectancy. Our larval studies suggest that elevated cytosolic calcium is present long before muscular activity has a chance to damage actin fibers or mitochondria. Future work will focus on assessing the role of calcium trafficking proteins in the cytosol of dystrophic muscles. This will allow us to identify candidate proteins that may be responsible for the initial increase in calcium that appear to kick off the chain of events leading to loss of mobility, morbidity, and death.

**Figure 6.**
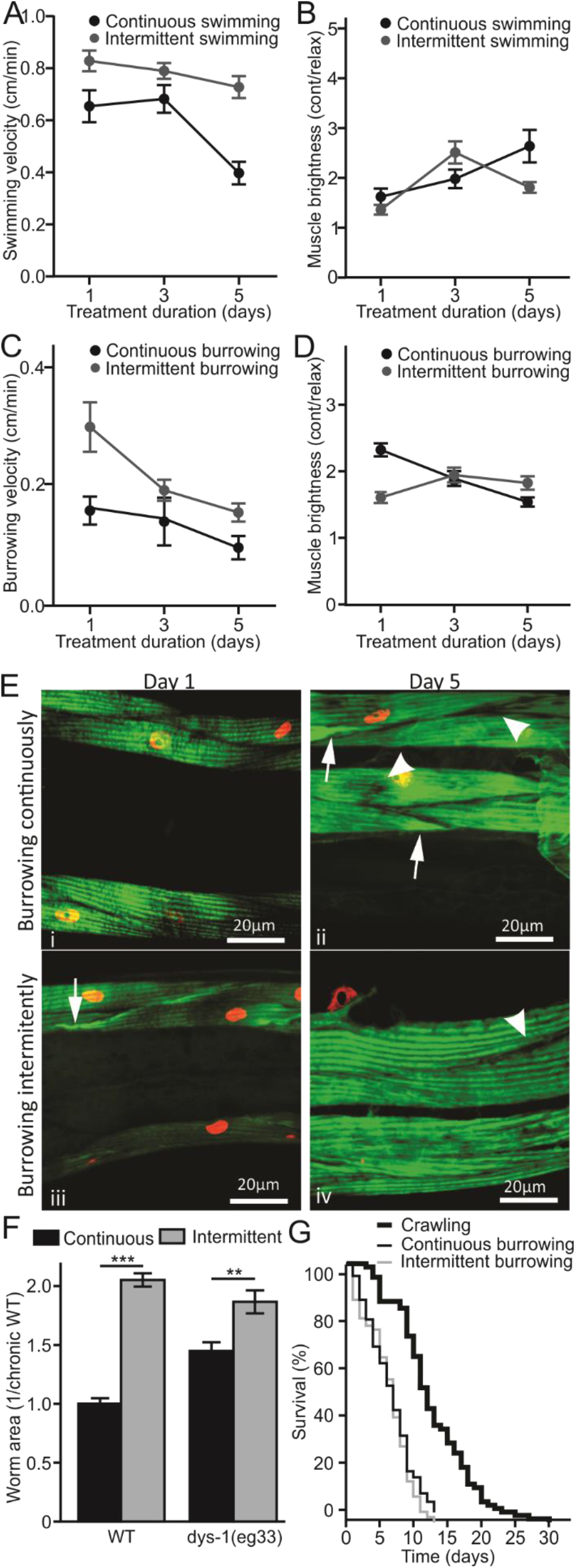
Intermittent exercise significantly improved some dystrophic muscle metrics, without affecting animal longevity. **A.** Dystrophic animals that swam daily for 90 minutes displayed a modest decrease in velocity over time. In contrast, animals that swam continuously had a significant drop in swimming velocity after day 3. **B.** The amplitude of calcium transients was initially similar for dystrophic worms swimming either continuously or intermittently. In both treatment calcium transients increased significantly over time, with continuously swimming animals seeing the greatest change. **C.** Worms that burrowed daily for 90 minutes had the greatest velocity after one exercise session. However, this effect disappeared by day 3. In both (intermittent and continuous) conditions velocity declined over time. **D.** While animals that burrowed continuously had a steady decline in the amplitude of cytosolic transients during locomotion, animals that burrowed intermittently saw a significant increase in these transients following their first burrowing session. **E.** Worms that burrowed chronically for five days showed signs of degeneration (arrowheads), as well as actin buildup in the attachment plaques (arrows). Intermittent burrowing animals displayed these signs of degeneration, but to a lesser extent. **F.** After 5 days, animals that burrowed intermittently were larger than those that burrowed continuously in both *dys-1* and WT strains, although the difference was smaller in dystrophic animals. **G.** Despite differences in velocity, calcium transients, and muscular integrity, animals that burrowed intermittently showed no improvement in longevity over those that burrowed continuously (Cox proportional hazard p = 0.194). ***p<0.001, **p<0.01, *p<0.05.

## ACKNOWLEDGEM ENTS

Funding provided by NIH NIAMS award (1R15AR068583-01A1), and by Illinois State University Startup funds and Research Grant (URG) to AV-G. We would like to thank Drs. Hongkyun Kim and Jon Pierce for strains, Dr. Kevin Edwards for assistance with confocal imaging, as well as Chance Bainbridge, Yazmine Giliana, Sruthi Singaraju, and Lavanya Sathyamurthy for contributing to some of the experiments presented here. Some strains were provided by the CGC, which is funded by NIH Office of Research Infrastructure Programs (P40 OD010440).

---

**Supplementary figure 1.**
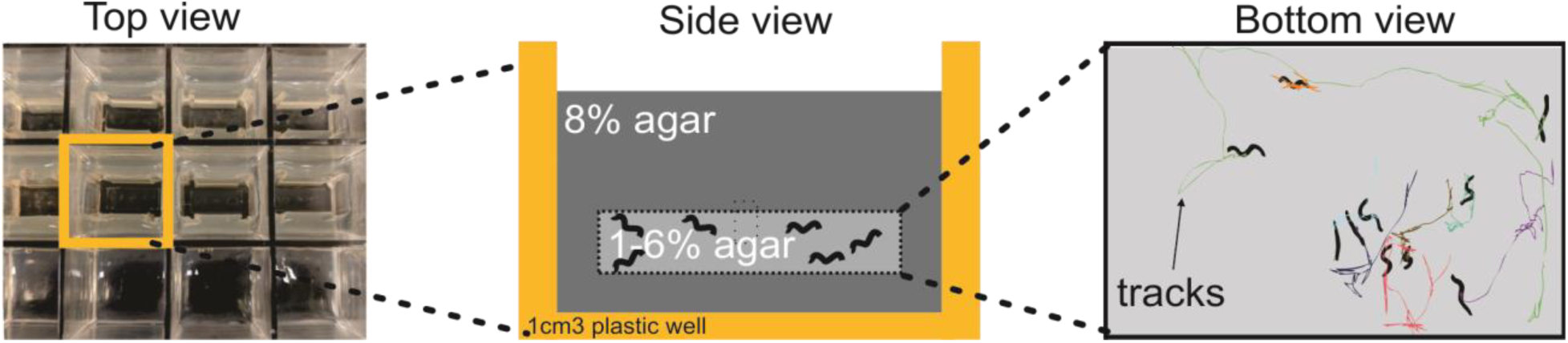
Assessment of burrowing animals. To facilitate the study of burrowing animals, we created an assay that allowed us to run several consecutive replicates of animals burrowing under different conditions. The schematic shows portion of a 100 well plate (left panel). Each well contained 8% agar which worms find difficult to burrow through. Within the 8% agar, agar of test densities (ranging from 1 to 6%) was placed after mixing in bacterial worm food (OP50). Worms were then injected into this test volume and covered n 8% agar to restrain their locomotion to the test volume. The burrowing layer was made thin (~1mm) to reduce vertical translation and aid in filming the animals. Burrowing worms were then filmed, and had their centroids tracked using Image-Pro-7 (Media Cybernetics, Rockville MD, USA) to obtain their burrowing velocity. The use of a fluorescent light source further allowed us to film and evaluate muscular exertion in animals expressing GCaMP2 in their musculature. Only animals burrowing in the horizontal plane were analyzed (right plate).

**Supplementary figure 2.**
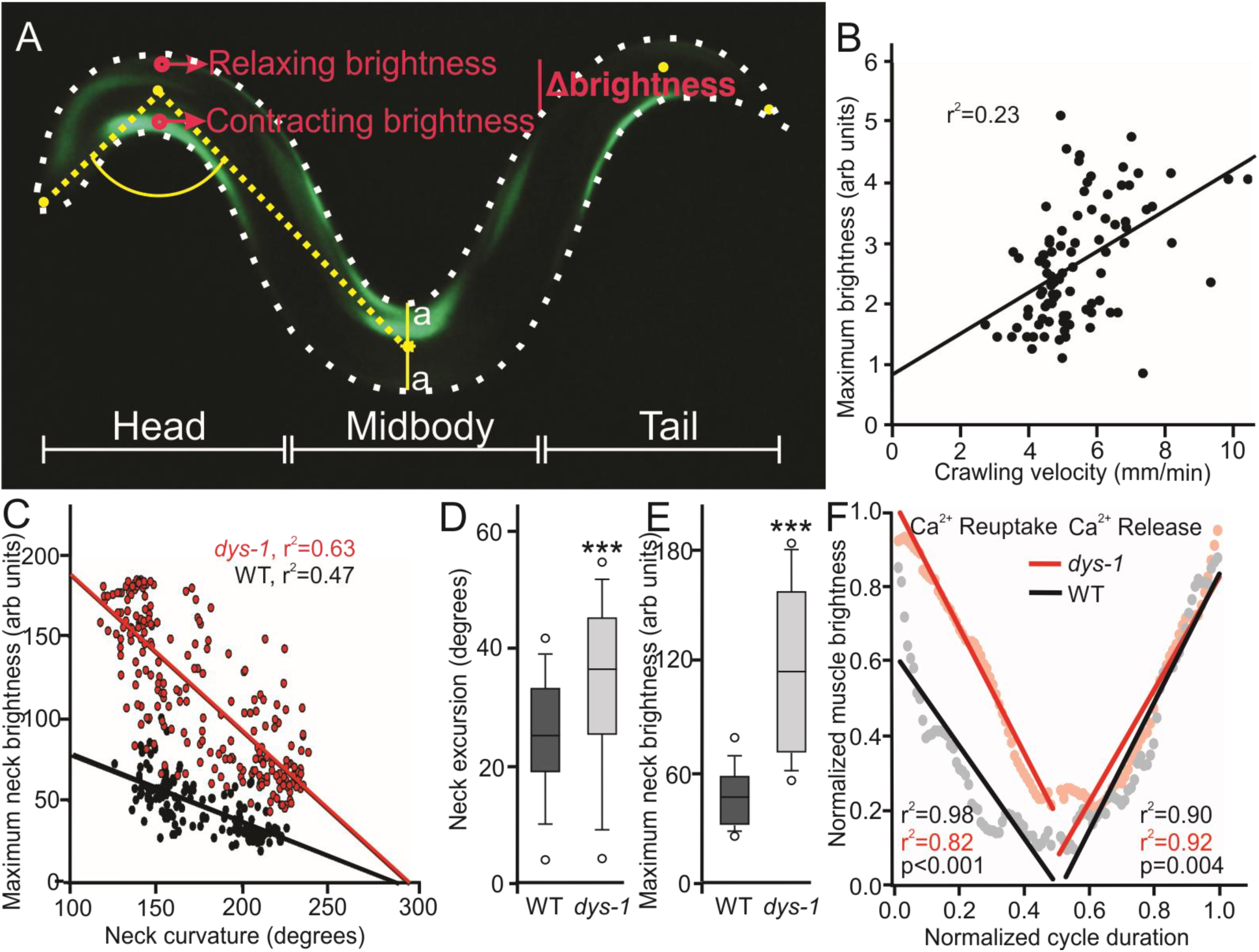
**A.** Schematic representing our method for measuring GCaMP2 brightness in the muscles of moving animals. Animals moving in the plane of the camera were selected. Measurements were obtained from the dorsal and ventral muscles immediately posterior to the pharynx (head), posterior to the vulva (midbody), and anterior to the anus (tail). ImageJ was used to measure body angle at selected segments, and to measure the brightness of antagonist body wall muscles. Measurements were either made at the apex of body curvature (to obtain maximal brightness during contraction), or over the entire cycle of movement (to measure calcium kinetics over time). **B.** Maximal animal brightness correlated with animal velocity (r^2^=0.23), with faster animals producing brighter (calcium) signals. **C.** Calcium transients also correlated with contraction amplitude, with greater brightness resulting in greater changes in body curvature for wild-type (r^2^=0.47) and dystrophic animals (r^2^=0.63). **D.** Dystrophic animals produced significantly larger neck excursions during locomotion (deeper bends), which were accompanied by greater calcium signals as measured by GCaMP2 **(E). F.** Comparison of intracellular calcium levels during contraction. During the relaxation phase of the cycle (0.0-0.5), when cytosolic calcium is actively pumped into the sarcoplasmic reticulum, we found that dystrophic worms differed significantly from wild-types in having an increased rate of removal (steeper slope), but also in failing to clear the cytosolic calcium before the beginning of the next contraction (x-intercept>0.5). Both wild-type and dystrophic strains had similar rates of calcium release (and same y-intercepts), although dystrophic worms were significantly slower than wild-types.

**Supplementary figure 3.**
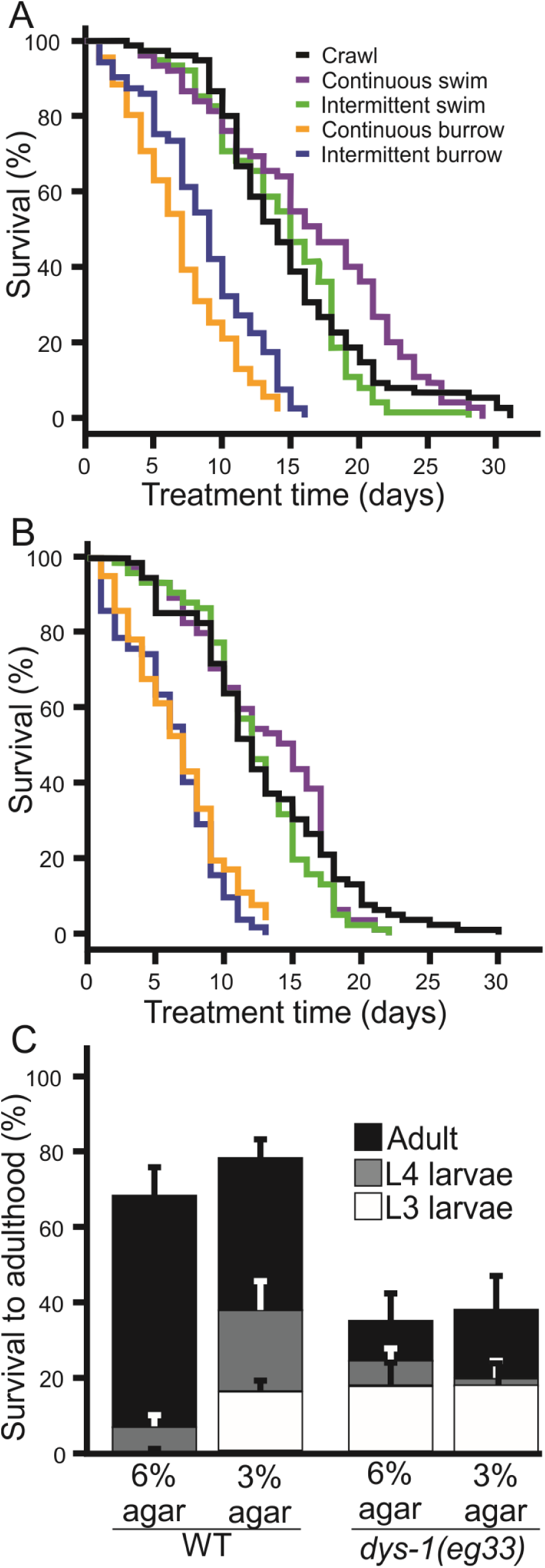
Intermittent exercise did not significantly increase longevity in dystrophic animals. **A.** Effect of the different exercise treatments used in this study on the longevity of healthy and dystrophic animals (**B**). Animals burrowing continuously had the shortest life span for both wild-type and dystrophic strains. However, while burrowing daily for 90 minutes (followed by crawling for 22.5hrs) had a reduced effect in the longevity of wild-type worms, this attenuation was absent in dystrophic animals. Dystrophic worm longevity was determined by the amplitude of exercise and not affected by its frequency. This effect was also observed for swimming animals. Dystrophic worms swimming intermittently had similar longevity to worms crawling continuously (which requires greater exertion).

**Supplementary figure 4.**
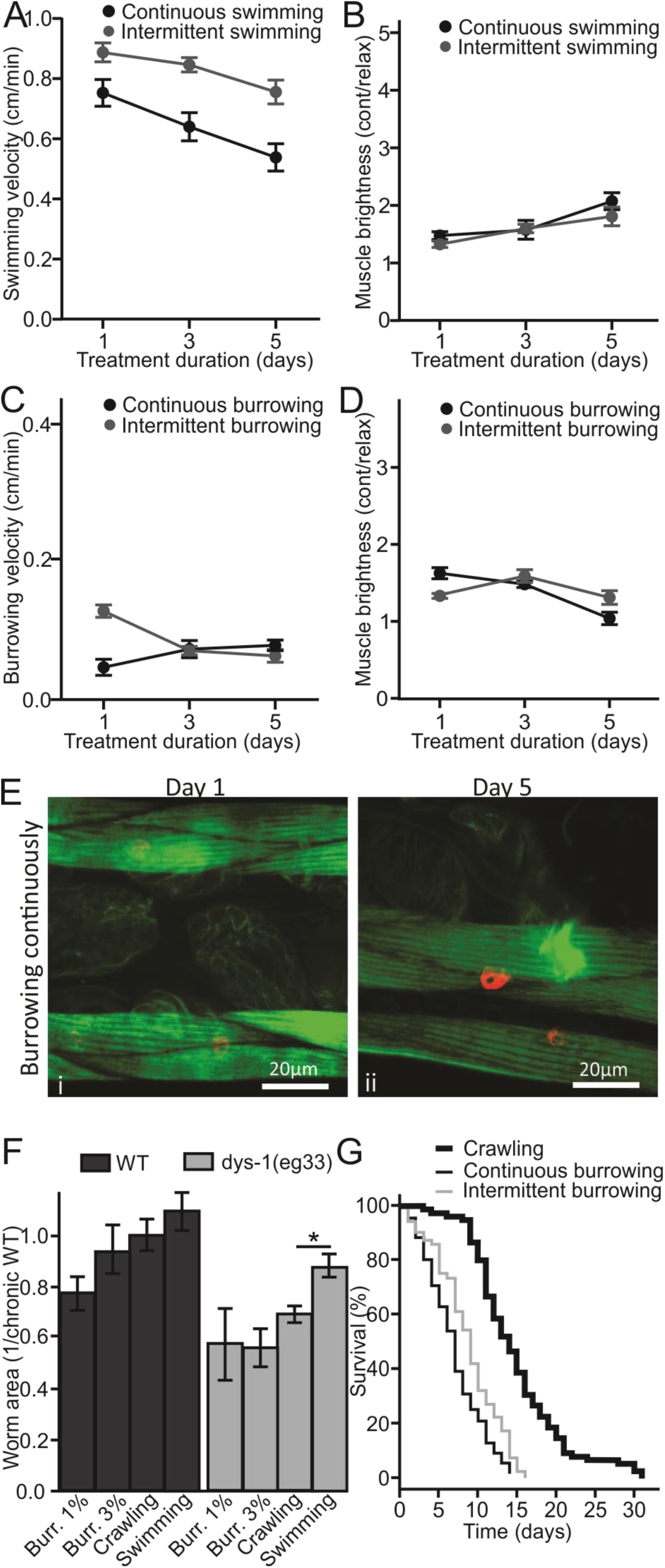
Wild type *C. elegans* longevity depended on the extent and duration of exertion. **A.** Animals that swam daily for 90 minutes displayed a similar decrease in velocity over time as animals that swam continuously, although intermittently swimming worms had consistently higher locomotion. **B.** The amplitude of calcium transients was similar for dystrophic worms swimming either continuously or intermittently across five days. In both treatment calcium transients increased significantly over time. **C.** Similarly, to dystrophic animals, worms that burrowed daily for 90 minutes had the greatest velocity after one exercise session and declined to match continuously-burrowing animals by day 3. **D.** While animals that burrowed continuously had a steady decline in the amplitude of cytosolic transients during locomotion, animals that burrowed intermittently saw an increase following their first burrowing session that declined by day 5. **E.** Worms that burrowed chronically for five days showed no signs of actin buildup in the attachment plaques **F.** After 5 days of continuous treatment, swimming dystrophic animals showed significant increase in size over crawling animals. **G.** Similar to differences in velocity, calcium transients, and worm size, wild-type animals that burrowed intermittently had slightly improved longevity over those that burrowed continuously (Cox proportional hazard **p = 0.00599).

